# A Single-Cell Atlas of the Mouse Dural Meninges Reveals Pervasive Sex Differences Across Cellular Compartments

**DOI:** 10.64898/2026.05.12.721894

**Authors:** Nikhita Arun, Natalie M. Frederick, Gabriel A Tavares, Ashleigh Vicchiarelli, Dheeraj Swaminathan, Jennifer Powers, Dimitrios Davalos, Cornelia C. Bergmann, Antoine Louveau

## Abstract

The dural compartment of the meninges forms a dynamic interface between the brain and the periphery, hosting diverse immune, vascular, mural and fibroblast populations. Single-cell studies have begun charting meningeal cellular diversity, yet a comprehensive view encompassing all major cellular compartments, intercellular communication, and the influence of sex remains lacking. Here, we present a single-cell transcriptomic atlas of the adult mouse dural meninges, profiling all major cell types in male and female mice at steady state. We uncover broad sex differences in cell-type-specific proportion, transcriptional programs, intercellular communication, and disease relevant signatures. Histological and cytometry analyses validate the biological relevance of these findings, establishing this atlas as a foundation for studying meningeal contributions to neurological disease in a sex-aware manner.

## Background

The dural meninges occupy a unique position at the interface between the central nervous system and the periphery^1–4^. With its fenestrated vasculature^5^, direct communication with the skull bone marrow^6–9^, and an extensive lymphatic network that drains cerebrospinal fluid from the central nervous system (CNS)^10–13^, the dura is simultaneously exposed to local and systemic signals, and capable of influencing CNS homeostasis, neuroinflammation and disease progression^14–28^. Over the past decade, studies have established that meningeal immune cells, including T cells^18,19,29,30^, mast cells^27,31^ and innate lymphoid cells^32–34^, can shape normal behaviors and inflammatory responses. The meningeal compartment has since been implicated in the pathology of numerous neurological disorders spanning neurodevelopmental, neuroinflammatory and neurodegenerative pathologies^14–28^.

Single-cell transcriptomic approaches have begun to reveal the cellular complexity underlying these meningeal functions^6,20,24,35–44^. Several atlases have been generated, primarily focusing on particular cell subtypes present in the meninges such as fibroblasts^35,39^, vascular cells^20^, and immune cells^6,37,40^. However, published datasets to date have been generated from single or mixed sex samples without disaggregation. However, virtually all neurological diseases with meningeal relevance show marked sex differences in prevalence, severity, or outcome, including but not limited to Alzheimer’s disease, meningitis, multiple sclerosis, traumatic brain injury or autism spectrum disorder^45–49^.

Sex is a fundamental biological variable that shapes immune cell identity, vascular function, and stromal cell biology across tissues^50–55^. In the peripheral immune system, females generally mount stronger antigen presentation responses driving faster infection clearing to the detriment of increased risk of developing autoimmunity^51,52,56^. On the other and, males tend to have a heightened innate immune system with stronger immunosuppression driving an increased efficacy of cancer therapies^57,58^. Sex differences in vascular biology are equally well established, with sex hormones regulating endothelial cell function^54,59^, adhesion molecule expression^60^, and immune cell trafficking across tissues^54,61^. Whether these systemic sex differences are recapitulated or amplified within the specialized meningeal microenvironment remains entirely unknown.

Here, we present a sex-stratified single-cell transcriptomic atlas of the mouse dural meninges, profiling all major cell compartments, under physiological conditions, in adult male and female mice. We show that sex is a pervasive organizing principle of meningeal biology, shaping cellular composition, transcriptomic programs, intercellular communication, and disease-relevant gene signatures across every compartment examined. We further demonstrate that sex-enriched transcriptomic profiles in meningeal cells are concordant with genes implicated in sex-biased neurological diseases, establishing this atlas as a resource for investigating meningeal contribution to CNS disease in a sex-aware manner.

## Results

### Single cell RNA sequencing of the meningeal compartment of male and female mice

The meningeal tissue, and particularly the dural compartment, is a complex and rich structure composed primarily of fibroblasts^35,62^ and blood vessels^5^. The dura also harbors a large and diverse immune cell population^20,24,37,63^ that can affect brain function in both normal and pathological conditions^16,20,62,64^. The dural layer of the meninges, because of its unique location on the outer-most border of the central nervous system (CNS)^1,5,62^ also presents characteristics of peripheral tissues, with the presence of peripheral nerves and associated myelinating cells^65^. To better appreciate the complexity and diversity of cells composing the dura, we collected the skull adherent meninges (dura with limited arachnoid) from 5 young adult male and female mice.

Immune cells (CD45+) and non-immune cells (CD45-) were FACS sorted and sequenced using the 10x Genomics single cell platform (**Figure 1A**). Clustering analysis of all sequenced cells after filtering for low quality cells using the Bioturing platform confirmed the isolation of all types of cells present in the dura, namely immune (*Ptprc*), endothelial (*Pecam1*), fibroblasts (*Col1a2*), mural (*Pdgfrb*) and neuroglial (*Plp1*) (**Figures 1B and 1C**). We first confirmed the overlap of all the major cell clusters across the UMAP for both male and female samples (**Figure 1D**). Furthermore, we validated that cells isolated from the male mice expressed high level of Y chromosome associated genes *Uty* and *Ddx3y,* while female cells expressed high levels of X chromosome associated genes *Xist* and *Tsix* (**Figure 1E**). We found similar proportions of the different cell types in both male and female samples suggesting proper isolation and representation (**Figure 1F**). Pseudo-bulk differentially expressed gene (DEGs) analysis on all cells unsurprisingly demonstrates that the most differentially expressed genes between the male and female cells are sex-chromosome associated genes (**Figure 1G**). The top 7 most DEGs (*Xist*, *Tsix*, *Erdr1y*, *Uty*, *Ddx3y*, *Eif2s3y* and *Kdm5d*) were subsequently excluded from the DEGs analysis (**Supplementary Table 1**). DEGs analysis for the 4 main groups of cells present in the dura found 200-400 DEGs- both up and downregulated in fibroblasts, immune cells and endothelial cells, but mural cells with only around 50 DEGs (**Figure 1H; Supplementary Table 1**). Comparison of the DEGs between the different group of dural cells demonstrated that the majority of these DEGs are unique to the cell type with only 18 DEGs shared between the endothelial, immune and fibroblasts and 4 DEGs shared between the 4 major cell types (**Figure 1I**). These data overall demonstrate the dural compartment consists of a diverse community of cells and that sex differentially affects each of them.

**Figure 1:**
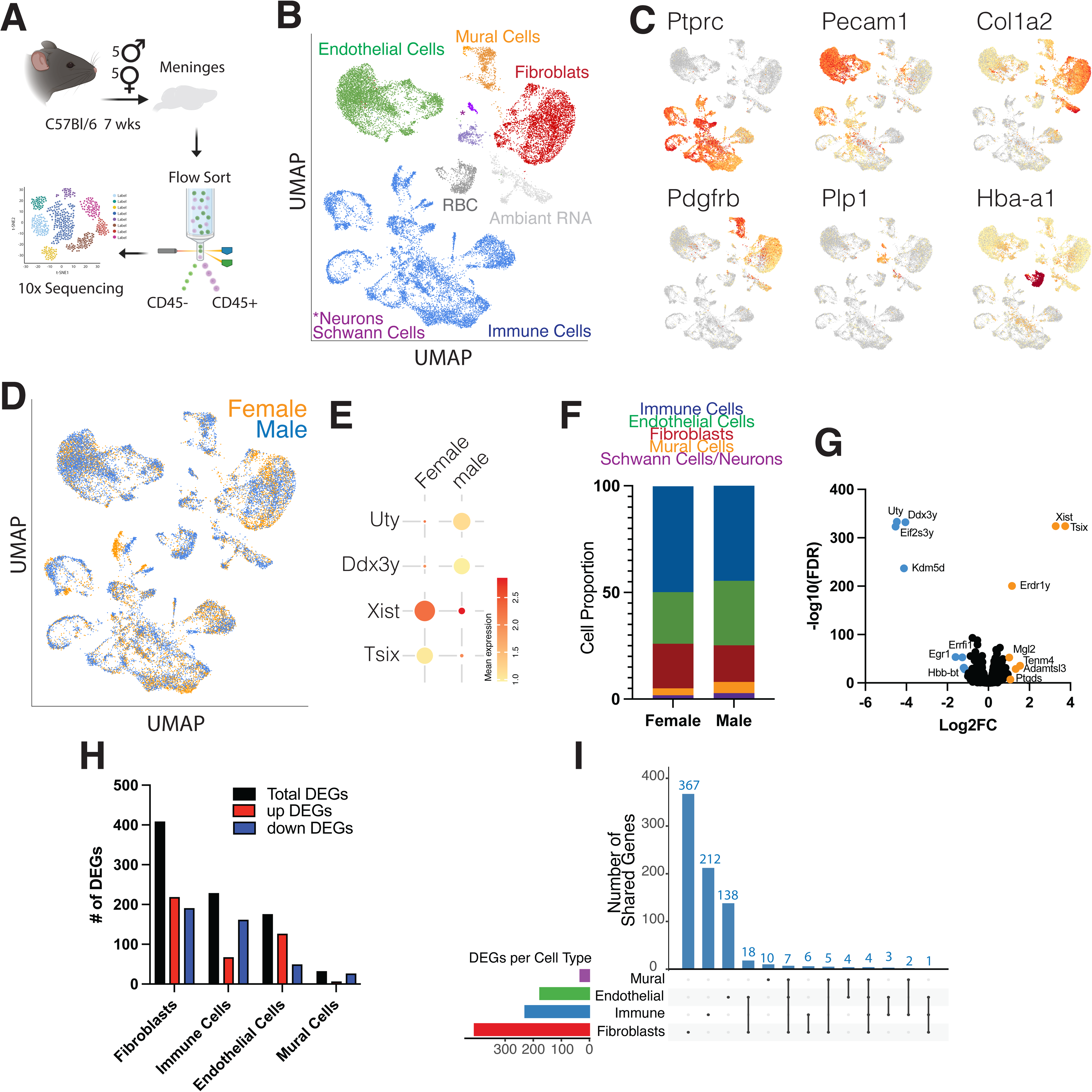
Single cell isolation and clustering of dural cells from male and female mice. A. Scheme of the experiment. Dura from male and female mice were isolated and pooled. CD45+ and CD45- cells were FACS sorted and analyzed using the 10x Chromium Genomic platform. B. UMAP clustering of the major cell types pooled from male and female mice. C. Feature plots of expression of the major markers identifying major dural cells types. D. UMAP clustering of the dural cells from male and female mice based on sex. E. Bubble plot of expression of sex-chromosome genes in male and female samples. F. Quantification of the proportion of major cell types in male and female samples. G. Violin plot of differentially expressed genes (DEG) from pseudo-bulk analysis comparing male and female cells. Colored genes depict significant differential expression. Blue is enriched in male sample; orange is enriched in female sample. H. Quantification of the number of DEG, either total, (black), up regulated (red) or down regulated (blue) is each major cell type comparing male and female mice. I. Upset plot comparing the unique and shared DEG between males and females across the major cell types.

### Sex impacts the composition of meningeal immune cells

Given the diversity of DEGs among the different cell types populating the dura, we isolated the immune cells and performed re-clustering (**Figure 2A**). In accordance to previously published data^37^, we found a large diversity of immune cells naturally present in the meninges (**Figure 2A**) that we identified using canonical, previously defined^37^, markers (**Figure 2B**). Separation of cells from male and female mice demonstrated similar clustering (**Figure 2C**) suggesting that the diversity of immune cells is similar in male and female mice. While there isn’t a category of immune cells that is absent in one sex, we next sought to investigate if some immune clusters may be more overrepresented in one sex. We compared the representative proportion of each immune subtype in male and female meninges **(Figure 2D**). We found that some immune cell types, mainly CD206+, CD209f+, MHCII+ and CD3e+ macrophages and ILC2s seem to be more abundant in female meninges while activated macrophages, IgM+ B cells, monocytes, neutrophils and gamma delta T cells are more prevalent in male mice (**Figure 2D**). Flow cytometry was used to confirm these sex-disparity observations. Like the prediction from the single cell analysis, we found that meningeal ILC2s are more abundant in female mice compared to male littermates (**Figures 2E and 2F**), a feature that is observed in other peripheral tissues^66^. IgM+ B cells but no other B cell subtypes appear to be more numerous in male meninges compared to female (**Figure S1A and S1B**). There were no differences in the number of monocytes or neutrophils (**Figures S1C and S1D**). Looking more specifically at the activation status of these cells, we similarly found no difference in the number of mature (CXCR2+) neutrophils (**Figure S1E and S1F**) or migrating (CCR2+) monocytes (**Figure S1G and S1H**). Single cell analysis highlights numerous subclusters of meningeal macrophages that represent both cell of different origin and function but also different activation status^21,37,37,44^. Flow cytometry markers are still currently unavailable to allow identification of all these subtypes. We identified 5 markers allowing the identification of all clusters. Lyve1 is upregulated in CD209f+ macrophages (**Figure 2B, Figure S1I**) and CD226 is particularly abundant in CD3e+ macrophages (**Figure S1I**). MHCII and CD206 (*Mrc1*) are highly expressed in the MHCII+ and CD206+ subclusters respectively but both are also high in migrating and activated macrophages (**Figure 2B, Figure S1I**). Using this combination of markers, we looked at the different subcluster of macrophages by flow cytometry. We found that the overall number of meningeal macrophages is higher in female mice compared to male mice, an increase driven by an increased number and proportion of MHCII+ and MHCII+/CD206+ (**Figure 2G,2H and S1J**). These results demonstrate that male and female meninges have different proportions of dural macrophage subtypes. We found no change in the number of αβ T cells, γδ T cells, NK and NKT cells (**Figure S1K and L**). We did however find an increase in the ratio of CD4/CD8 T cells in female mice compared to male (**Figure S1M**), a feature that is observed in blood and peripheral tissue^67^. Overall, these data demonstrate differences in immune cell composition, particularly macrophages, ILC2s and IgM+ B cells in male and female dura.

**Figure 2:**
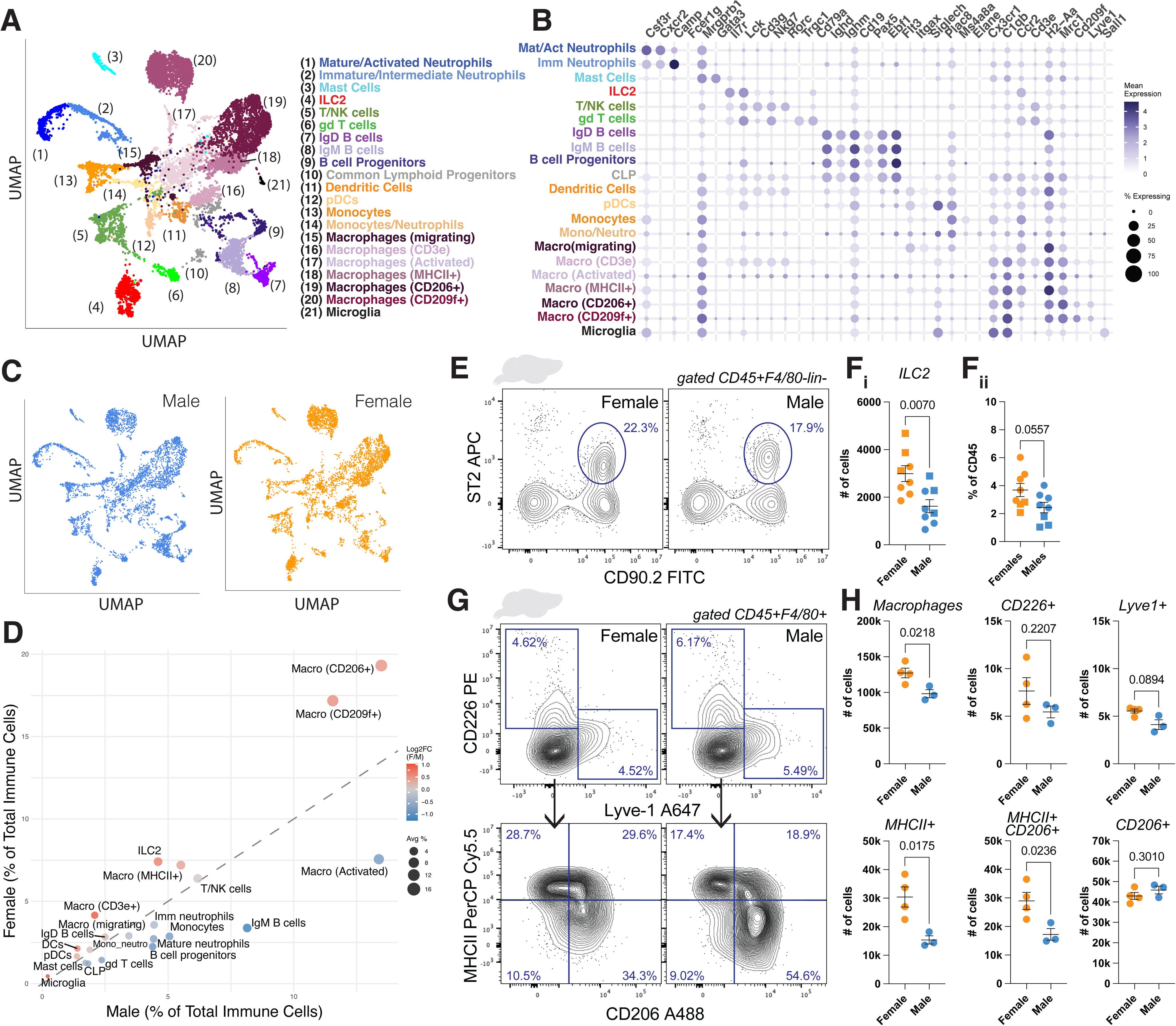
Sex affects the proportion of meningeal immune cells. A. UMAP of the dural immune cells (CD45+) from male and female mice. B. Bubble plot of expression of major immune markers identifying the identity of dural immune cells. C. UMAP of dural immune cells segregated according to sex. D. Cell proportion scatter plot of immune cell clusters between male and female mice. E. Representative contour plot of meningeal ILC2s in male and female mice. F. Quantification of the number (i) and percentage of CD45 (ii) ILC2s in the meninges of male and female mice. Mean ± sem. Welch’s test. G. Representative contour plot of meningeal macrophages in male and female mice. H. Quantification of the number of macrophages in the meninges of male and female mice. Mean ± sem. Welch’s test.

### Sex-disparity in the transcriptomic profile of dural ILC2s

We next investigated how sex affects the transcriptomic profile of dural immune cells. Analysis of the number of DEGs shows that ILC2s are the immune cells with the highest numbers of DEGs between males and females (**Figure 3A, Supplementary table 2**) followed by different subtypes of macrophages (**Figure 3A**). We therefore focused on understanding how sex affects the transcriptome of meningeal ILC2s. Independent clustering of the dural ILC2s shows almost no overlap between male and female cells (**Figure 3B**) illustrating the large number of DEGs between them (**Figure 2A**). Volcano plot mapping of the ILC2s DEGs between male and female mice reveal upregulation of genes related to calcium sensing and neuroendocrine and cytokine secretary programs *(Rbm3*, *Mapk12*, *Cadps2*, *Pcsk1*, *Mctp1*)^68–71^ in female cells, while genes related to reduced ILC2 function (*Calca*, *Nt5e*)^72–74^, and increased Nfκb signaling (*Nfkb1*)^75,76^ are observed in male cells (**Figure 3C, Supplementary table 2**). Using GSEA pathway enrichment analysis to extract how observed differences in gene expression might affect the activities of the meningeal ILC2s, we found that their sex-dependent transcriptomic profile impacts their function on many levels (**Figure 3D, Supplementary table 2**). Among altered functions are metabolic reprogramming and signaling and immune responses (**Figure 3D**). Dural ILC2s also appear to have a different response to type II cytokines and IL33 signaling (**Figure 3D**). Sex also appears to regulate the implantation of the dural ILC2s into their environment with alteration in proliferation, survival and migration (**Figure 3D**). Overall, these data suggest a profound effect of sex on the transcriptomic profile of meningeal ILC2s. To better understand how sex alters the function of meningeal ILC2s at the molecular level, we extracted the genes from the DEGs analysis that belonged to the different categories highlighted by the GSEA analysis (**Figure 3E**). We found an increase in *Klrg1* in male meningeal ILC2s (**Figure 3E**), suggesting that male ILC2s may be retained more persistently in the meningeal compartment^77,78^. This hypothesis is supported by the increased expression of integrins by male ILC2s, that could favor the anchoring of these cells to the extracellular matrix component of the meninges. Interestingly, male ILC2s exhibited increased expression of the gene encoding ST2 (*Il1rl1*), the IL33 receptor^79^ and also appear to have the transcriptional machinery to respond to ST2s engagement, with increased expression of the NFκB and JAK pathways^75,80^ (**Figure 3E**). On the contrary, female ILC2s have the molecular machinery ready for cytokine production with increased *Mapk12*^69^ and *Cadps2*^70^ expression(**Figures 3C and 3E**). Overall, the single cell transcriptomic analysis of meningeal ILC2s demonstrates that sex alters their tissue retention, proliferation capacity and response to cytokines. We used flow cytometry to confirm these transcriptomic observations at the protein level. Analysis of the level of expression of ST2 in meningeal ILC2s demonstrates increased expression in male ILC2s (**Figure 3F and 3G**), in agreement with the RNA sequencing data (**Figure 3E**). We next tested the response of dural ILC2s to their canonical cytokine IL33^79^. Meningeal ILC2s were isolated from the dura of male and female mice and stimulated with IL33 ex vivo for 6h. Female ILC2 presented an increased activation profile compared to male cells at 6h with upregulation of CD25 (**Figures 3D and 3I**). Simultaneously, we found a higher proportion of female meningeal ILC2s produce IL5 (**Figures 3J and 3K**). These results suggest that female ILC2s respond more prominently to IL33, potentially because of their increased expression of the molecular machinery that regulates activation and cytokine secretion.

**Figure 3:**
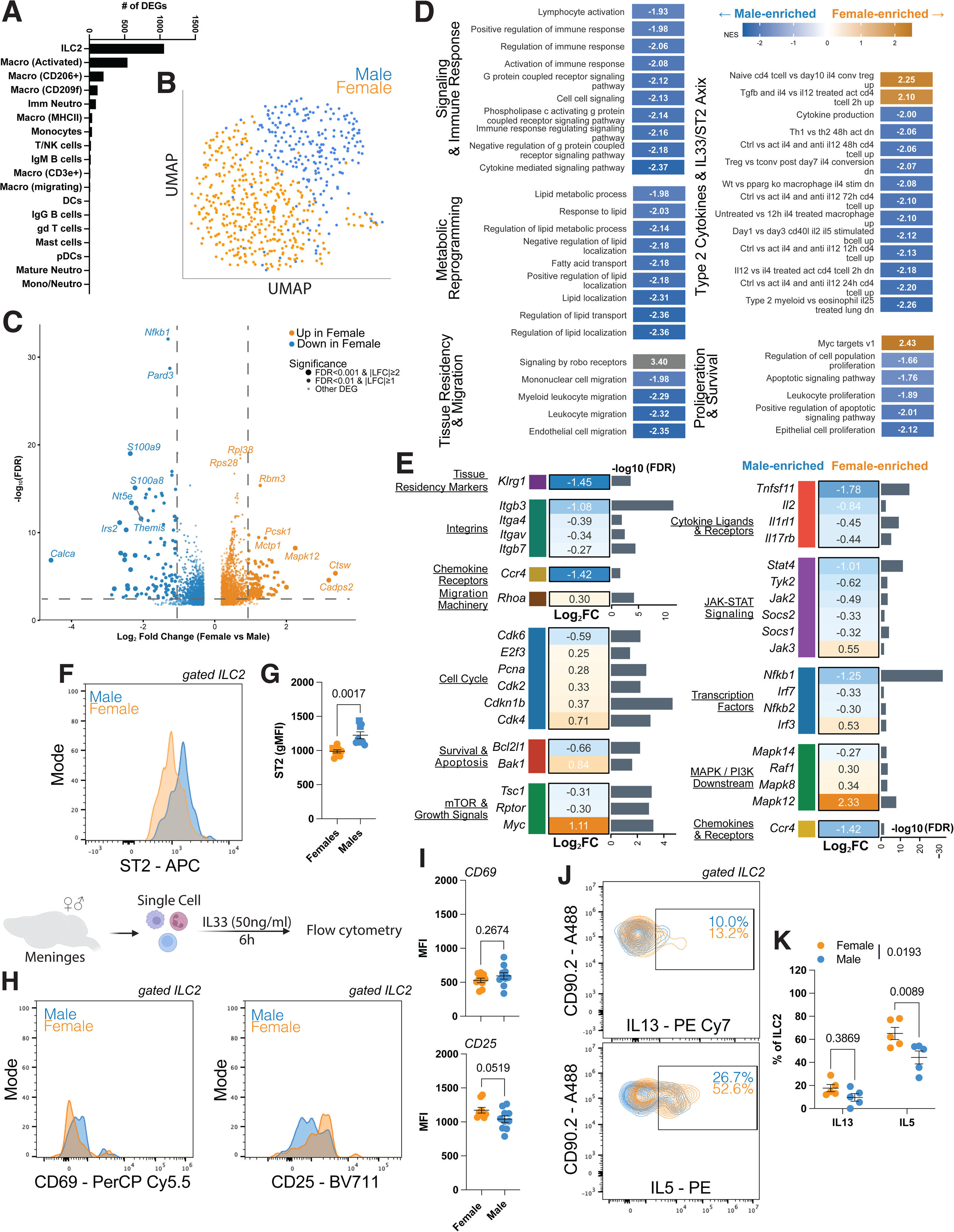
Dural ILC2s phenotype is modulated by sex. A. Quantification of the number of differentially expressed genes in meningeal immune cell clusters. B. UMAP clustering of meningeal ILC2s from male and female mice. C. Volcano plot of the differentially expressed genes in meningeal ILC2 between male and female mice. D. Bar plot of normalized enrichment scores from Gene Set Enrichment Analysis (GSEA) of pathways grouped by functional category between male and female dural ILC2s. E. Diverging bar plot of individual genes grouped by functional category comparing male and female dural ILC2s. F. Representative histogram of ST2 expression by male and female dural ILC2s. G. Quantification of the geometric mean fluorescent intensity of ST2 on male and female dural ILC2s. Mean ± sem. Welch’s test. H. Meningeal immune cells from male and female mice were isolated and stimulated ex-vivo with IL33 (ST2 ligand) for 6h. Representative histogram of expression of CD69 and CD25 by male and female dural ILC2. I. Quantification of the geometric mean fluorescent intensity of CD69 and CD25 expression on male and female dural ILC2s. Mean ± sem. Welch’s test. J. Representative contour plots of IL13 and IL5 expression by male and female meningeal ILC2s. K. Quantification of the percentage of cytokine producing dural ILC2s in male and female mice. Mean ± sem. Two-way repeated measure ANOVA with Sidak’s multiple comparisons test.

### Sex regulates the transcriptomic profile of the vascular compartment

Although broadly studied in immune cells, sex differences have also been reported in other cell types and several tissues throughout the body^81,82^. Therefore, we next investigated how sex may also regulate the non-immune cells comprising the meningeal compartment. Endothelial cells (EC) were clustered independently from other meningeal cell types. As previously demonstrated^20^, we found endothelial cells of the entire blood vascular tree (artery, capillary and veins), a population of blood endothelial cells (BEC) belonging to the arachnoid/pial layer, and a lymphatic endothelial cells (LEC) population (**Figure 4A**). All cells were identified using established cell identity markers^20,83^ for brain and meningeal vasculature (**Figure 4B**). Segregation of the cells based on sex demonstrated that all clusters are represented in both sexes (**Figure 4C**). To better understand the effect of sex on endothelial cells, we generated the list of DEGs for each endothelial cluster. We found that all clusters except for Bmx+ arterial ECs and LECs had 50 or more DEGs with capillary 2 and arachnoid/pial endothelial cells having over 150 DEGs (**Figure 4D, Supplementary Table 3**). These results suggest a profound effect of sex on the endothelial cells of the dura. We then extracted DEGs from the list based of their association with functional pathways essential for endothelial cells. We particularly looked at genes involved in cell adhesion, immune activation, migration and chemotaxis, and extracellular matrix interactions pathways. Comparison of the fold change and direction of these genes across the different meningeal EC clusters revealed specific patterns (**Figure 4E**). Female EC seem particularly enriched for integrins (*Itga2, Itga5, Itgav, Itgb1, Itgb4*) and integrin signaling (*Jam2, Jam3, Mcam, Alcam*) (**Figure 4E**), but also more immunologically active with overexpression of MHCII (*H2-Aa*. *H2-Ab1*, *H2-Eb1*) in venous, capillary and arterial ECs (**Figure 4E**). In contrast, male venous EC express higher levels of the immune adhesion markers *Icam1* and *Vcam1*, aiding immune cell extravasation and migration in male cells^84,85^ (**Figure 4E**).

**Figure 4:**
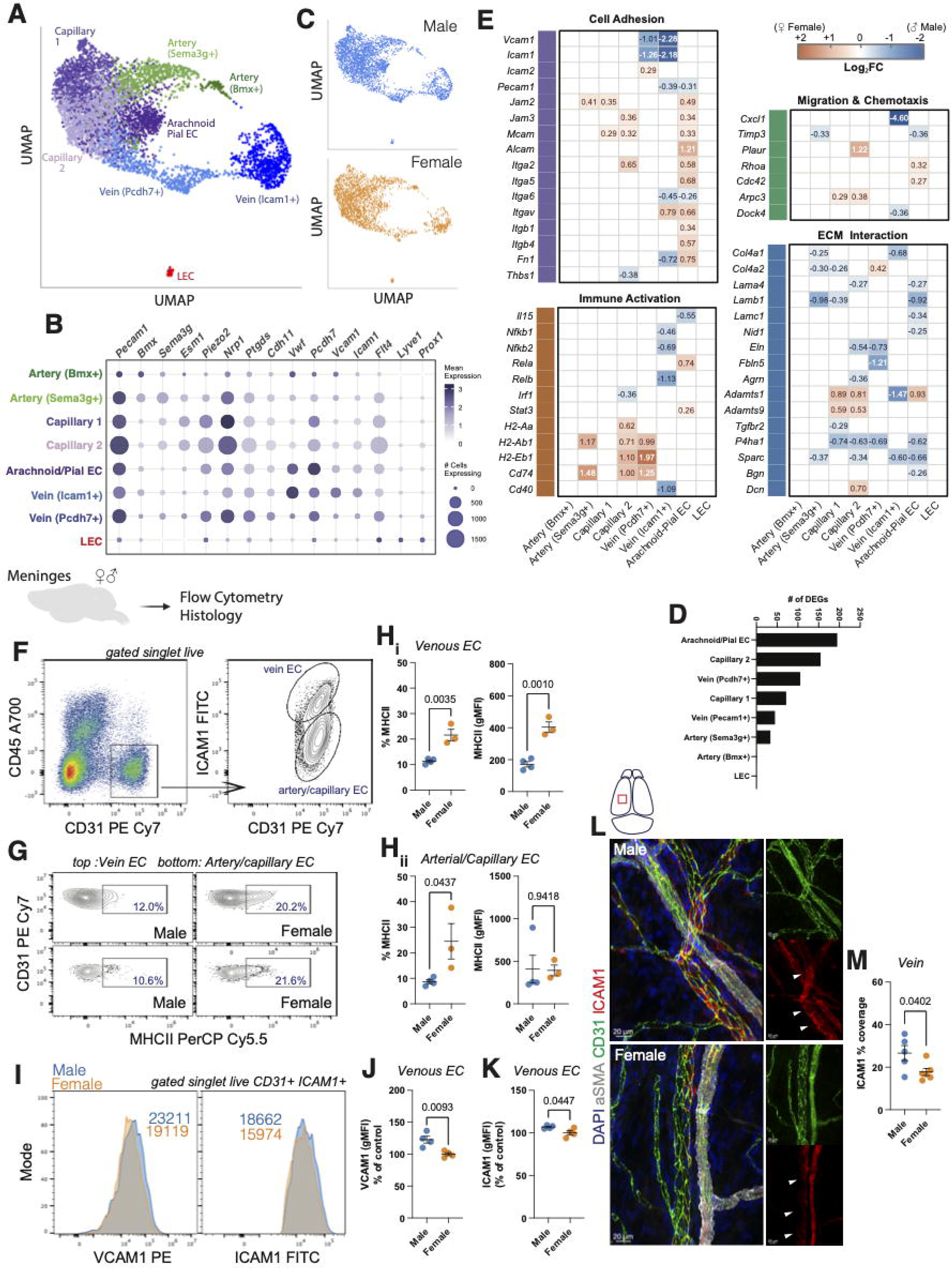
Sex influences the transcriptomic profile and phenotype of dural endothelial cells. A. UMAP clustering of dural endothelial cells of male and female mice. B. Bubble plot of expression of major endothelial markers identifying vascular subtypes in the dural compartment. C. UMAP clustering of vascular cells from male and female mice. D. Quantification of the number of differentially expressed genes in the dural endothelial clusters between male and female mice. E. Diverging Heatmap of genes organized into functional categories in the different vascular dural clusters comparing male and female mice. F. Representative dot and contour plot for identification of venous, arterial and capillary endothelial cells in the dura of mice. G. Representative contour plots of MCHII expression in dural endothelial subtype in male and female mice. H. Quantification of the percentage of cells expressing MHCII, and geometric mean fluorescence intensity of MHCII expression by dural endothelial subtype in male and female mice. Mean ± sem. Unpaired t test. I. Representative histogram of VCAM1 and ICAM1 expression by dural venous endothelial cells. J. Quantification of the geometric mean fluorescence intensity of VCAM1 expression by venous endothelial cells between male and female mice. Results are expressed as percentage of control (male condition). Mean ± sem. Unpaired t test. K. Quantification of the geometric mean fluorescence intensity of ICAM1 expression by vein endothelial cells between male and female mice. Results are expressed as percentage of control (male condition). Mean ± sem. Unpaired t test. L. Representative images of ICAM1 (red) expression by dural vasculature (CD31; green) in male and female mice. αSMA (grey) identifies arteries. M. Quantification of the ICAM1 percentage of coverage in venous endothelial cells of male and female mice. Mean ± sem. Unpaired t test.

We confirmed some of these predictions using flow cytometry and histology. A previous report demonstrated that Icam1 is a marker for venous EC of the dural, particularly sinus endothelial cells^20^. We identified brain endothelial cells by the expression of CD31 and lack of CD45 expression (**Figure 4F**). We identified venous EC as ICAM1+ while capillary and arterial cells were ICAM1 negative (**Figure 4F**). In accordance with the single cell RNAseq prediction, we found that female venous EC (**Figures 4G and 4H**), and to a lesser extent arterial/capillary cells (**Figures 4G and 4H**) have a higher expression of MHCII, and higher proportion of MHCII+ cells compared to male cells.

Analysis of venous expression of ICAM1 and VCAM1 showed a mild but significant upregulation of both adhesion marker in male ECs (**Figures 4J -4K**). Given the mild difference and the reliance on ICAM1 as a venous EC marker, we further examined its expression by histology. We focused our analysis on veins (characterized by the lack of smooth muscle cell coverage and large diameter) away from the sinuses, to avoid confounding ICAM and VCAM1 signal from surrounding fibroblasts and smooth muscle cells. Quantification of ICAM1 coverage in dural vein confirmed significantly higher expression of ICAM1 in male compared to female littermate (**Figure 4L and 4M**). Our data demonstrate that sex impacts the transcriptomic profile of the dural endothelial cells and predicts biological differences potentially relevant for the function of the meningeal compartment.

### Sex regulates the transcriptomic profile of the fibroblast compartment

The third major cluster of cells in the mouse dura comprises fibroblasts. Sub-clustering of the fibroblasts highlighted the diversity of cells with identification of 7 different fibroblasts subclusters (**Figure 5A**). Two of them, periosteal and suture-associated express genes related to bone biology indicative of the outer dural layer that directly attaches to the skull^39,62,86^ (**Figure 5A**). We identified two clusters forming the inner dura membrane^39^. We observed more specialized and unique clusters such as the osteogenic fibroblasts that express some markers associated with bone biology (*Runx2*^87^), but also myofibroblast markers (*Tnc*^88^, *Lrcc15*^89^), perivascular fibroblasts enriched for vascular marker (*Flt1*^90^) and arachnoid fibroblasts (*Ptgds*^39^) (**Figure 5A-C**). Separation of fibroblasts based on sex showed the presence of all fibroblast clusters in both sexes (**Figure 5B**). Quantification of the number of DEGs in dural fibroblasts revealed that the outer dura fibroblasts are the most affected by sex, followed by inner dura fibroblasts 2 (**Figure 5D, Supplementary Table 4**). To extract more biologically significant information relating to the fibroblasts function in male and female mice, we extracted relevant functional genes. We found that male fibroblasts, across the spectrum are enriched for genes related to the immediate early response, ER stress and proteostasis (**Figure 5E**). The major divergence between male and female fibroblasts appears to be related to the extracellular matrix. Male fibroblasts express higher level of ECM genes such as *Col1a1*, *Col1a2*, *Col4a1*, while female fibroblasts have an enhanced anti-fibrotic signature (**Figure 5E**). Interestingly, we also found that male inner dura fibroblasts have increased expression of IL33, a cytokine signaling through ST2 on ILC2s^79^ (**Figure 5E**).

**Figure 5:**
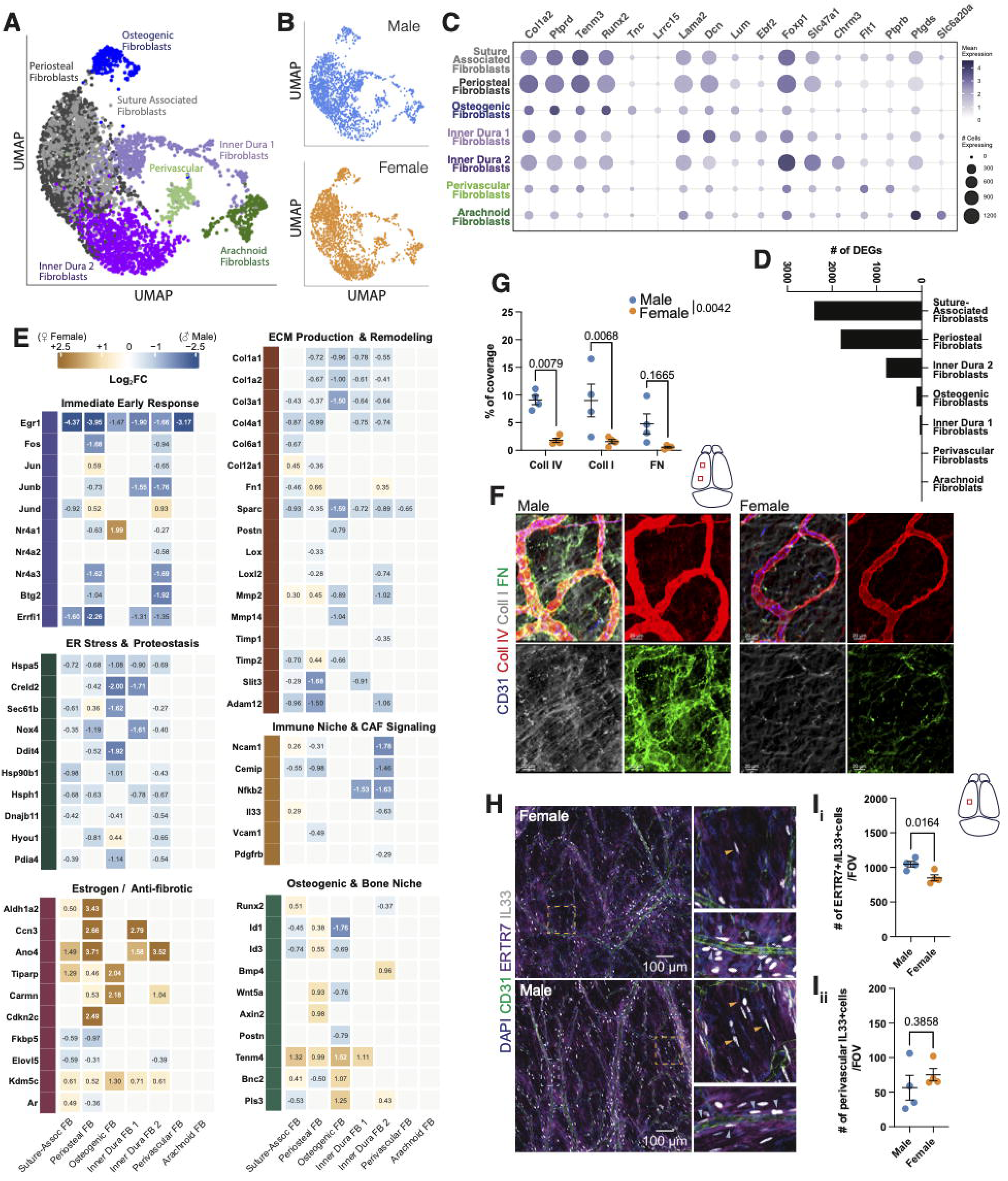
Dural fibroblasts transcriptome and phenotype is regulated by sex. A. UMAP clustering of dural fibroblasts populations of male and female mice. B. UMAP clustering of dural fibroblasts segregated by sex. C. Bubble plot of expression of major fibroblasts markers identifying the dural fibroblasts clusters. D. Quantification of the number of differentially expressed genes in the dural fibroblasts clusters between male and female mice. E. Diverging Heatmap of genes organized into functional categories in the different dural fibroblasts clusters comparing male and female mice. F. Representative images of extracellular matrix deposition in the meninges of male and female mice. G. Quantification of the coverage of extracellular matrix component by dural fibroblasts in male and female mice. Mean ± sem. Two-way repeated measure ANOVA with Sidak’s multiple comparisons test. H. Representative images of IL33 (grey) expression by dura fibroblasts (ERTR7, purple) and perivascular cells (CD31, green) in male and female mice. I. Quantification of the number of IL33+ fibroblast (i) and perivascular cells (ii) per field of view in male and female mice. Mean ± sem. Unpaired t test.

To confirm these observations at the protein level, we first immunolabelled dura from male and female mice for extracellular matrix component (**Figure 5F**). We focused the analysis on the extracellular matrix outside of the blood vasculature and its basement membrane to be specific to the looser fibroblastic extracellular matrix^91^ (**Figure 5F**).

Quantification of the coverage of collagen I, IV and fibronectin deposition on the dural fibroblasts showed increased coverage in male compared to female mice (**Figure 5F and 5G**), reflecting the prediction of the RNA sequencing. Given the strong difference in ILC2s number and phenotype, we further explored the expression of IL33 in the dural meninges. We found that the number of fibroblasts (ERTR7+) expressing IL33 is higher in male compared to female mice (**Figure 5H and 5I**). The quantification was performed away from the sinus to exclude the suture-associated fibroblasts. Interestingly, IL33 is also expressed by endothelial/perivascular cells in the meninges^92^. We found that, contrary to the fibroblasts, the number of perivascular IL33+ cells is not different between sexes (**Figure 5I**). Overall, these data demonstrate that our single cell RNA sequencing accurately predicts the effect of sex on the biology of meningeal fibroblasts. It also demonstrates that sex influences the transcriptome of dural fibroblasts, suggesting that sex likely modulates the biological response of dural fibroblasts.

### Sex influences cell-cell interactions in the dura

The dura is a complex structure, within which cells are constantly interacting and influencing each other’s function. We next wondered if sex could not only affect the transcriptome of the dural cells but also influence their cell-cell interaction. We therefore performed CellChat analysis^93,94^ on our dural data. Suture-associated and periosteal fibroblasts were excluded from analysis because of their location, which limits their interaction with the immune and vascular cells. LEC and arterial (Bmx+) cells were also excluded because they did not meet minimum cell number per cluster threshold. Our analysis focused on the differences of cell-cell interactions between male and female mice rather than looking at how cells interact in the dura. Comparison of the number of statistically significant interactions between the different dural cell types between male and female mice demonstrates important sex-driven disparities (**Figure 6A, Supplementary Table 5**). Mast cells and inner dura fibroblasts 2 demonstrate an overall increased number of interactions with dural cells in female mice (**Figure 6A**), while Lamc3 pericytes and smooth muscle cells have increased numbers of interactions in male mice (**Figure 6A**). This suggests that sex will influence interactions within the dural environment. In accordance with the fact that ILC2s have the highest number of differentially expressed genes between male and female mice (**Figure 3A**), we found significant alterations in cell-cell interactions involving ILC2s (**Figure 6B and 6C**). Using CellChat analysis, we compared the interactions between male and female conditions based on the differential expression of ligand-receptors and the probability of interaction occurrence. We found that male ILC2s appear to interact with the meningeal vasculature via the TGFβ pathway, while the interaction in female mice is primarily driven by PECAM1-PECAM1 interaction (**Figure 6B**). Both TGFβ and PECAM1 signaling regulate vascular tone and immune cell circulation^95–99^, suggesting that male and female ILC2s may regulate the meningeal vasculature differently. We then compared the maximum probability of interaction of ILC2s with each dural cell cluster for each ligand-receptor pair between male and female mice. We found that female ILC2s have a higher probability of interaction with meningeal macrophage populations and endothelial cells compared to male ILC2s (**Figure 6C**), demonstrating that sex significantly alters the cellular interactome of ILC2s.

**Figure 6:**
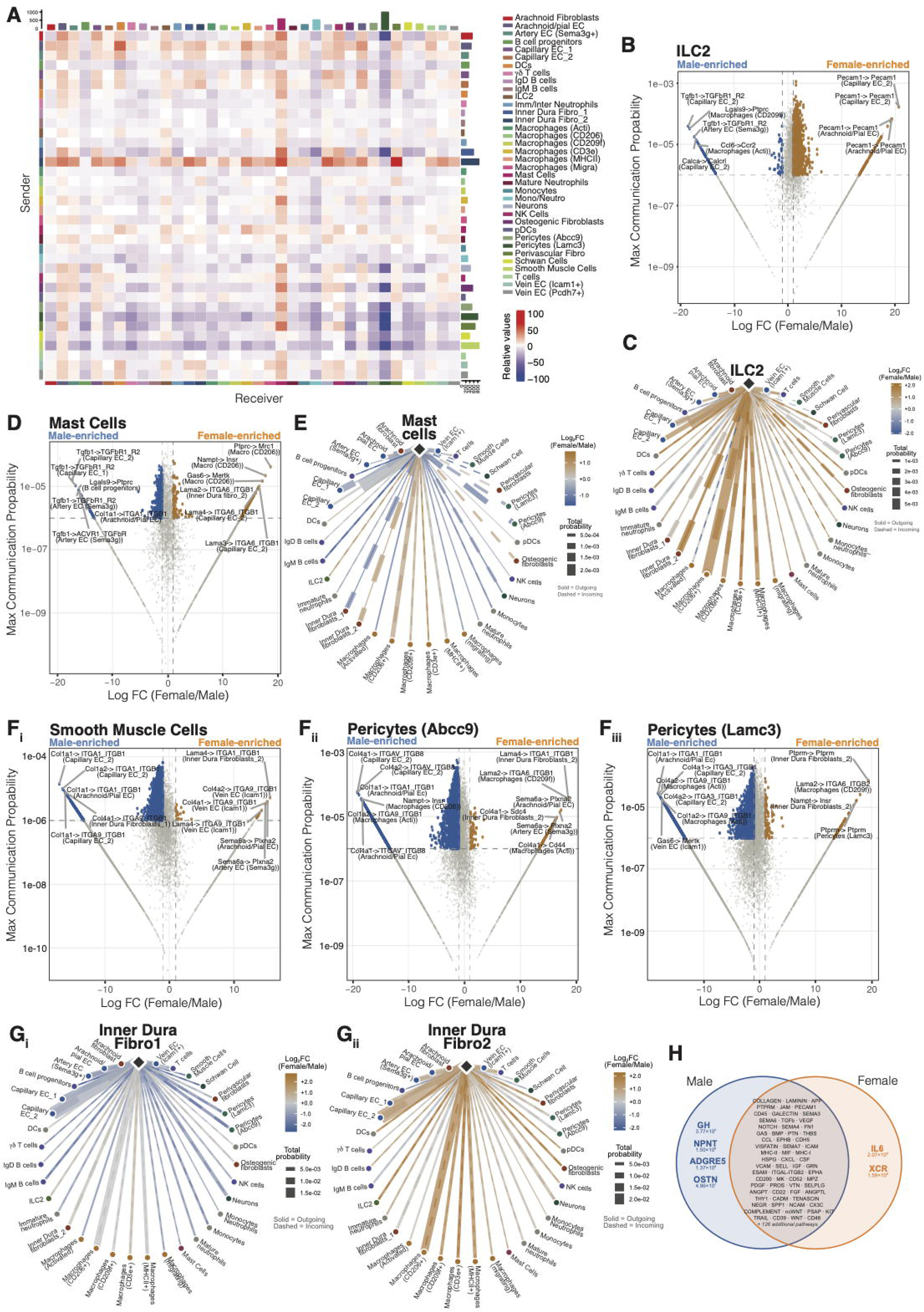
Sex influences the interactome of dural cells. A. Heatmap of number of significant interactions between male and female dural cells. B. Scatter plot of differential ligand-receptor pair usage for ILC2s between male and female mice. C. Chord diagram of sex-differential communication probability of dural ILC2s with other dural cells. D. Scatter plot of differential ligand-receptor pair usage for mast cells between male and female mice. E. Chord diagram of sex-differential communication probability of dural mast cells with other dural cells. F. Scatter plot of differential ligand-receptor pair usage for smooth muscle cells (i), Abcc9 pericytes (ii) and Lamc3 pericytes (iii) between male and female mice. G. Chord diagram of sex-differential communication probability of inner dura fibroblasts 1 (i) or 2 (ii) with other dural cells. H. Venn diagram of ligand-receptor pathway usage by male and female mice independently of cell identity.

We next investigated if cells that show little to no DEGs between male and female mice would still present changes in their interaction pattern. We focused on mast cells, as they showed no statistically significant DEGs between male and female mice (**Figure 3A**). Mast cells indeed presented a sex-biased interactome. At the molecular level, we found that male mast cells had enriched interactions with the meningeal vasculature using the TGFβ pathway, while female mast cells interacted with the meningeal vasculature using extracellular matrix component such as *Lama2*, *Lama3*, and *Lama4* (**Figure 6D**). Given the essential roles of mast cells in regulating the vasculature in the context of migraine^100^, these data support that sex may drive the different molecular pathways that are used. Furthermore male mast cells demonstrated a higher probability of interaction with vascular and mural cells and female mast cell with most macrophages’ subtypes (**Figure 6E**), suggesting that sex affects the biology of a cell beyond its cell intrinsic transcriptome.

We next explored the non-immune cells with high change in interactions between sex. Mural cells showed a significant increase in the number of interactions in male mice (**Figure 6A**). Analysis of the differential interaction pair probability showed that dural mural cells use different pathway to interact with the meningeal vasculature. In male mice, mural cells appear to use Collagen signaling including *Col1a1*, *Col1a2*, *Col4a1*, and *Col4a2* to interact with cells of the vasculature tree (**Figure 6F**). In female mice, other constituents of the extracellular matrix like laminins (*Lama4*, *Lama2*) and transmembrane receptors (*Sema6a*) appear to control the interaction with vascular cells (**Figure 6F**). Overall, this demonstrates that sex may change or rather refine how cells interact with each other, potentially to drive different or similar function.

We also found that cells of similar lineage have different cellular interactomes. Inner dura fibroblasts whether belonging to group 1 or group 2 show different probability of interaction. Inner dura fibroblasts 1 show a higher probability of interaction with the majority of dural cells in male mice, while inner dura fibroblasts 2 have higher interaction probability in female mice (**Figure 6G**). Given that we did not see a change in the proportion of these cells in the male and female dura (**Figure 5B**), these results suggest that sex may influence dural fibroblasts are the preferred interacting partner to other dural cells. Investigating ligand-receptor pair usage in male and female mice, regardless of cell type involved, demonstrated that some pathways (Growth Hormone, Nephronectin, Adhesion G Protein Coupled Receptor E5 and Osteocrin) are uniquely used by male cells, while IL6 and XC Chemokine Receptor are uniquely used by female cells (**Figure 6H**). The sexual dimorphism of the Growth Hormone pathway is not surprising given its direct regulation by sex^101^. The other pathways are, however, more surprising and highlight sex as a likely central regulator of dural biological regulation.

Collectively, these data demonstrate that sex shapes dural cell function beyond directly transcriptomic regulation, and underscore the value of this atlas as a powerful tool for dissecting sex-dependent differences in meningeal biology.

### Sex disparity in cell specific gene expression correlates with sex disparity in neurological disorders

Most, if not all, neurological diseases present strong sex-disparities with Alzheimer’s disease^102^, multiple sclerosis^103^, Lupus^104^, depression^105^ and migraine^106^ affecting preferentially females while glioblastoma^107^, autism spectrum disorders^108^, Parkinson’s disease^109^, fragile X^110^ or meningitis^111^ tends to affect males predominantly. We thus interrogated how the sex specific gene enrichment in the dural cells may correlate to the known lists of genes that are involved in the sex-disparity of the previous neurological disorders. We found a high degree of concordance between the genes enriched in endothelial cells and fibroblasts and the genes regulating sex disparity (**Figure 7A, Supplementary Table 6**). Particularly, there is a very strong concordance in the male fibroblast’s genes and the male prevalent diseases, suggesting that male fibroblasts may influence male prevalent neurological disorders (**Figure 7A**). To our surprise, we also found a more ambiguous correlation of sex-bias genes in dural immune cell types with both concordant, discordant and mixed correlations, suggesting that the potential implication of the dural immune cells in the preponderance of sex-biased neurological diseases is likely more complex (**Figure 7A**). We then examined specific genes, and how, in different group of dural cells, they correlate with male or female prevalent neurological disorders. We found that *Ccn3*, *Fos*, *H2-Eb1* and *Cd74* are enriched in female endothelial cells and in female biased neurological disease while *Cxcl1* is particularly enriched in male EC and correlates with male-prevalent disease, particularly meningitis (**Figure 7B**). We similarly found that female fibroblasts gene correlate with female biased diseases and some male fibroblasts genes correlate with male biased diseases (**Figure 7C**). Among immune cells, only macrophages show some degree of correlation (**Figure 7D**), albeit at lower levels than endothelial cells and fibroblasts, while lymphoid cells show limited correlation (**Figure 7E**). Overall, these results support that the sex-enriched genes expression in specific dural cell populations correlate with genes pathways underlying neurological disease disparity.

**Figure 7:**
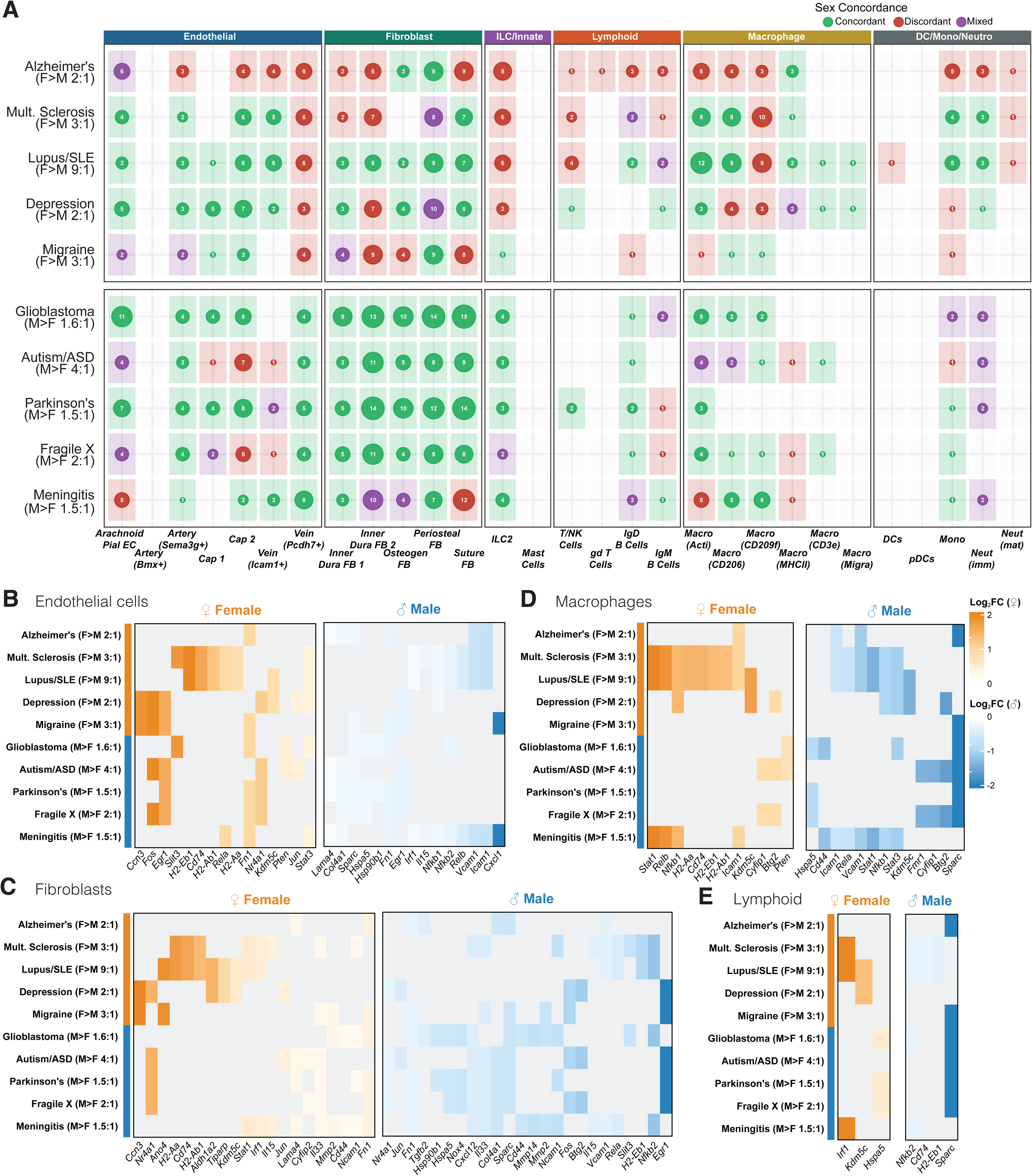
Correlation of sex-specific gene signatures of dural cells with disease associated gene enriched pathways. A. Dot matrix of overlap between sex-biased differentially expressed genes in dural cell types and genes associated with neurological and autoimmune diseases with documented sex prevalence differences. B. Split diverging heatmaps of disease associated sex-biased DEGs in dural endothelial cells. C. Split diverging heatmaps of disease associated sex-biased DEGs in dural fibroblasts. D. Split diverging heatmaps of disease associated sex-biased DEGs in dural macrophages. E. Split diverging heatmaps of disease associated sex-biased DEGs in dural lymphoid cells.

## Discussion

The dural meninges occupy a unique immunological position, simultaneously interfacing the system circulation through the fenestrated vasculature^5^, sampling CSF from the brain parenchyma^112^, and receiving immune input from the adjacent skull bone marrow^6,7^. The convergence of peripheral and central immune signals makes the dura a critical gatekeeper of CNS homeostasis^62^, yet a comprehensive, sex-aware characterization of its cellular composition has been lacking. Here we present a single- cell transcriptomic atlas of the mouse dural meninges encompassing all major cellular compartments in both sexes and demonstrate that sex is a pervasive organizing principle of meningeal cell identity, shaping cell composition, transcriptomic programs, intercellular communication, and disease-relevant gene signatures across every cell compartment examined.

Meningeal ILC2s emerged as one of the most profoundly affected cell type affected by sex in the dura, exhibiting differences in abundance, transcriptional identity, and functional responsiveness. Recently, meningeal ILC2s have been implicated in the formation of inhibitory synapses during normal development through IL-13 production^32^, an essential process for normal social behavior. Female meningeal ILC2s are more abundant and demonstrate greater cytokine secretion capacity upon IL-33 stimulation, including higher IL-5 production and upregulation of the secretory machinery genes *Cadps2* ad *Mctp1*. This may confer a form of developmental resilience in female mice. If the meningeal ILC2s to interneuron signaling axis operates above a threshold necessary for adequate inhibitory synapses formation, females may retain sufficient ILC2s function when this pathway is partially disrupted by genetic, environmental or inflammatory insults. Males, who’s baseline meningeal ILC2s abundance and cytokine output is lower, may already operate closer to this threshold, such that the same magnitude of disruption pushes the signal below the level required for normal inhibitory synaptogenesis. Neurodevelopmental disorders characterized by social behavior impairment, such as ASD and ADHD, affects males at approximately three to four times the rate of females^113^. Sex disparity in meningeal ILC2s may represent one of the underlying mechanisms that contribute to this clinical observation. Beyond development, meningeal ILC2s have been implicated in the immune response to injury, both in traumatic brain injury and ischemia^33,34^. In both instances, the release if IL33 by the injury site is a trigger of ILC2s activation^33,34^. Our data reveal that male and female ILC2s are differentially configured. Male ILC2s express higher levels of ST2 but respond less prominently to IL33 stimulation than female ILC2s. This apparent paradox may reflect differences in downstream signaling with apparent upregulation of the *Mapk12* and other components of the cytokine secretory machinery amplifying the signal transduction downstream of ST2 engagement. Whether this translates into differential meningeal ILC2s responses to CNS injury in male and female mice, and what the functional consequences for recovery might be, remains an important open question.

One broader theme emerges when comparing the transcriptional landscapes of multiple meningeal cell types across sexes. Female meninges trend toward more constitutive immune surveillance and antigen presentation, evidenced by MHCII upregulation across vascular endothelial cells and macrophage populations, a greater abundance of MHCII expressing macrophages, and ILC2s primed for rapid cytokine secretion. Male meninges, by contrast, trends towards immune cell mobilization and tissue retention, with higher ICAM1 and VCAM1 expression on venous endothelial cells, potentially facilitating leucocyte extravasation from the circulation^84,85^, integrin upregulation and *Klrg1* expression in ILC2s promoting tissue anchoring^77,78^, and increased ECM deposition by fibroblasts potentially remodeling the stromal scaffold for immune cell retention. These findings suggests that sex shapes not only the baseline immune state of the meninges, but also the preparedness of this tissue to respond to insults. The female meningeal environment, with its higher antigen presentation tone, may favor ongoing adaptive immune surveillance and containment of chronic immune threats, while the male environment, with its enhanced capacity for rapid cellular mobilization from the circulation, may favor a more reactive response to acute insults. Whether these baseline differences translate into sex-specific susceptibility or resilience across different neuroinflammatory conditions remains to be investigated. This atlas nevertheless provides a framework for addressing these questions experimentally.

The potential functional relevance of this atlas is reinforced by our correlation analysis, which reveals the intersection of sex-specific DEGs in meningeal cells with genes implicated in sex-biased neurological diseases. The upregulation of ICAM1 and VCAM1 in male dural endothelium, for example, may lower the threshold for immune cell entry into the meningeal space during neuroinflammatory challenge^114–116^, potentially contributing to worse outcomes in conditions such as bacterial meningitis, where male patients show higher susceptibility and mortality^111^. Among all cell types, male-enriched DEGs in dural fibroblasts show particularly strong concordance with genes associated with male prevalent disorders. This finding points to an underappreciated potential role for the meningeal fibroblasts compartment in mediating sex-biased disease susceptibility. Fibroblasts are not passive structural cells, they actively regulate ECM composition^117,118^, can produce cytokines and chemokines^119,120^, and engage in bidirectional signaling with the immune and vascular compartment^118,119^. The sex-specific transcriptional identity of these cells may therefore influence the meningeal microenvironment in ways that differentially predispose male and female tissue to specific pathological trajectories. Our data identify this as a largely unexplored direction for future investigation.

Several limitations of this study merit consideration. First, while we document extensive sex differences across the meningeal compartment, the upstream mechanisms driving these differences remain unresolved. Sex chromosome encoded gene regulation and circulating sex hormone levels are both plausible drivers and likely interact. Second, our atlas was generated at steady state in young adult mice. It is yet unknown whether the sex differences we describe are maintained, amplified, or diminished under disease conditions, or with aging.

## Conclusion

We present a sex-stratified single-cell transcriptomic atlas of the mouse dura meninges, demonstrating that sex is a central determinant of meningeal cellular composition, transcriptional identity, and intercellular communication across all major compartments. Female and male meninges exhibit distinct immunological, vascular and fibroblastic configurations at steady state with direct implications for how each sex may respond on neuroinflammatory challenges. This atlas provides a fundamental resource for the field and underscores the necessity of incorporating sex as a biological variable in meningeal and neurological research.

## Methods

### Mice

Mice were maintained and bred in-house under standard housing conditions (14/10h light dark cycles and fed ad libitum). All mice used were C57Bl/6j (JAX:000664) at 7 to 9 weeks of age. Experiments used littermates. Animals from different cages in the same experimental groups were selected to ensure randomization. Experimenters were blind to experimental groups during analysis. Sample sizes were determined based on previously published experiments. All experiments were approved by the Institutional Animal Care and Use Committee of the Cleveland Clinic Research.

### Tissue collection and processing

Mice were euthanized by intraperitoneal injection of Beuthanasia (100mg/kg) and perfused with 0.1M of PBS (10-12ml/mouse). For staining of extracellular matrix component (ECM), mice were subsequently perfused with PBS with 1% paraformaldehyde (PFA; Electron Microscopy Sciences; 15714). Skin was removed from the head, and muscles were stripped from the skull. After removal of the mandibles, the skull cap was dissected. For histology, the skull cap was fixed with PBS with 1% PFA for 2 to 24h at 4C (except for the ECM samples that were directly put in PBS). After fixation, the skull caps were washed with PBS and preserved in PBS/0.02% azide until staining. For flow cytometry, the skull caps were kept in FACS buffer until dissection.

### Immunohistochemistry

Whole mount meninges were incubated with PBS containing 2% of normal serum (Goat: Abcam; ab7481; Chicken: Jackson ImmunoResearch; 003-000-120), 1% bovine serum albumin (BSA; Equitech Bio; BAH70), anti-CD16/32 (1/1000; BioXCell; BE0307); 0.1% Triton-X-100 (Fisher Scientific; BP151-100) and 0.05% Tween 20 (Fisher Scientific; BP337-100) for 1h at room temperature (RT).

Subsequently, the tissues were incubated with appropriate dilutions of primary antibodies O/N at 4C in PBS containing 1% BSA and 0.5% Triton-X-100. Primary antibodies used are anti-alpha smooth muscle actin-Cy3 (1:200; Sigma Aldrich; C6198); Hamster anti-ICAM1 (1:100; Thermo Scientific; MA5405); Rabbit-anti-CD31 (1:200; Thermo Scientific; MA5-37858); goat-anti-Collagen I (1:200; SouthernBiotech; 1310-01); Rabbit anti-Collagen IV (1:200; BioRad; 2150-1470); sheep anti-fibronectin (1:200; BioRad; VPA00045); rat anti-CD31 (1:200; BioLegend; 102401); goat anti-IL33 (1:200, R&D System; AF3626) and anti-ERTR7 (1:200; Thermo Scientific; MA1-40076). Whole mounts were then washed 3 times for 5 minutes at RT with PBS followed by incubation with fluorescently conjugated or biotinylated secondary antibodies, or conjugated streptavidin (Jackson ImmunoResearch; Thermo Fisher Scientific) at 1/500 for 2h at RT in PBS with 1% BSA and 0.5% Triton-X-100. After 5 min incubation with DAPI (1µg/ml; Thermo Fisher; D1306), the whole mounts were washed 3 times for 5 min with PBS, dura were dissected from the skull cap and mounted with Aqua-Mount (Fisher Scientific; 14-390-5) on glass slides (Fisher Scientific; 12-550-15).

### Image analysis

Images were acquired with a Leica Stellaris confocal system (Leica Microsystems) using LAS X software (Leica Microsystems). Images were acquired with a 40x objective (NA 1.10 or 1.30) (Stellaris) with a resolution of 1024×1024 pixels with a z-step of 2 to 4 microns. Quantitative assessments were performed using FIJI software (NIH) or IMARIS (OXFORD Instruments). ICAMI expression levels was measured by the thresholded area of expression on vein (Vein were manually traced (large vessel without aSMA staining). Coverage of extracellular matrix components were measured via thresholding. The areas were manually selected to eliminate any signal coming from the vascular basement membrane. The density of IL33+ cells were manually counted per field of view. Deciphering there fibroblastic versus perivascular status was based of ERTR7 expression and localization around blood vessels.

### Flow cytometry

Meninges were dissected and digested for 15min at 37C with 1.4U/ml of Collagenase VIII (Sigma Aldrich; C2139) and 35U/ml of DNAse I (Sigma Aldrich; DN25) in complete media (DMEM (Gibco) with 2% FBS (Invitrogen), 1% L-glutamine (Gibco); 1% penicillin/streptomycin (Gibco); 1% sodium pyruvate (Gibco), 1% non-essential amino acid (Gibco); 1% essential amino acid (Gibco), and 1.5% HEPES (Gibco). The cell pellets were resuspended in ice-cold FACS buffer (0.1M PBS; 1mM EDTA (Fisher Scientific) and 1% BSA (Equitech-Bio; BAH70); pH 7.4). For IL33 stimulation, cells were stimulated with IL33 (50ng/ml) in complete media in the presence of Brefeldin A (1µl/ml;BD; 347688) for 6 hours. Cells were stained for extracellular markers with antibodies at a 1:200 dilution with 1:100 dilution of anti-CD16/32 (BioXCell; BE0307). Cells were stained for 30 min at 4C, then fixed in 1% PFA in 0.1M PBS. For intracellular staining, cells were fixed and permeabilized using the Cytofix/Cytoperm fixation and permeabilization kit (BD; 554714) following the manufacturer’s instructions. The following antibodies were used: anti-CD45 A700 (BD; 560510); anti-F4/80 PE cf594 (BD; 565613); anti-ST2 APC (BD; 567590), anti-CD90.2 FITC (Biolegend; 105316); anti-CD226 PE (Biolegend; 133603); anti-Lyve1 A647 (R&D System; FAB2125R); anti MHCII PerCP Cy5.5 (ThermoFisher; A12902); anti-CD206 A488 (BioRad; MCA2235A488); anti-CD69 PerCP Cy5.5 (Biolegend; 104521); anti-CD25 BV711 (Biolegend; 102049); anti IL13-PE Cy7 (Biolegend; 159407); anti-IL5 PE (Biolegend; 504303); anti-CD31 PE CY7 (Biolegend; 102418); anti-VCAM1 PE (Biolegend; 105713); anti-ICAM1 FITC (Biolegend; 116105); anti-B220 APC Cy7 (BD; 55204); anti-CD19 BV711 (BD; 563157); anti-IgD BV510 (Biolegend; 405723); anti-IgM PerCP Cy5.5 (Biolegend; 406511); anti-CD11b PE Cy7 (Biolegend; 101215); anti-Ly6G BV711 (Biolegend; 127643); anti-Ly6C BV570 (Biolegend; 128030); anti-CXCR2 APC Cy7 (Biolegend; 149313); anti-CCR2 APC (Biolegend; 150627); anti-CD90.2 APC (BD; 553007); anti-TCRgd PE Cy5 (Thermo Fisher; 15-5711-82); anti-TCRb BV711 (BD; 563135); anti-CD8a V450 (BD; 560471) and anti-CD4 PE Cy7 (BD; 552775). For ILC2s, lineage staining (BD; 561301) was used to exclude other immune cells. Fluorescence data were collected with a Cytoflex (Beckman Coulter), then analyzed using FlowJo 10 software (BD Life Sciences). Single cells were gating using the area and height of the forward scatter and the side scatter, then selected using the LIVE/DEAD Fixable Dead Cell Stain Kit per the manufacturer’s instruction (Invitrogen; L34980). An aliquot of unstained cells was counted using a Countess II cell counted (Thermo Fisher) using trypan blue to provide a cell counts.

### Single cell preparation

The cell preparation for the single cell RNA sequencing is the same as for regular flow cytometry with the addition of translation inhibitors (Actinomycin D (5µg/ml; Sigma Aldrich; A1410); Tripolide (10µM; Sigma Aldrich; T3652); Anisomycin (27.1µg/ml; Sigma Aldrich; A9789) to limit generation of digestion mediated artefact. Cells were stained for Calcein AM (1:1000; Invitrogen; C1430) to identify nucleated cells; DAPI (1/10000; ThermoFisher; D1306) to identify living cells and CD45 (1:200; ThermoFisher; 559864) to decipher CD45+ and CD45- cells. Cells were sorted using a FACS ARIA (BD) into BSA coated Eppendorf. CD45+ and CD45-from male and female mice were processed using the 10x Genomic Chromium platform (10x Genomics) following manufacturer’s instructions. Samples were sequenced using the NovaSeq at a depth of 50,000 reads per cell.

### Single cell analysis

Raw sequences reads were aligned to the mouse reference genome and gene expression matrices were generated using Cell Ranger (10x Genomics). Cell Ranger filtered genes were imported into BioTuring BBrowser for downstream analysis. Quality control filtering was applied to remove low-quality cells based on minimum gene count thresholds, maximum gene count thresholds (to exclude potential doublets), and percentage of reads mapping to mitochondrial genes. Genes detected in fewer than a minimum number of cells were also excluded from downstream analysis. Data from multiple samples were integrated to correct for batch effects using BioTuring’s built in integration. Normalized and scaled gene expression were used for principal component analysis (PCA), and the top principal components were used to construct a shared nearest neighbor (SNN) graph for unsupervised clustering. Cell clusters were visualized using Uniform Manifold Approximation and Projection (UMAP). Cell type identities were manually assigned to each cluster based on the expression of established marker genes for known dural cell population. Differentially expressed genes (DEGs) between male and female mice were identified within each cell type cluster using BioTuring BBrowwer. DEGs were called using Venice statistical test and filtered based on log2 fold-change (above 0.5) and false discovery rate (inferior to 0.01). The number of statistically significant DEGs per cluster was used to assess transcriptional sex differences across dural populations. Cell type proportion were calculated with BioTuring BBrowser. Visualization of gene expression, cluster markers, and proportion was performed using ggplot2 custom R scripts.

To identify biological pathways enriched in a sex-biased manner across dural cell population, two complementary approaches were applied in R. Gene Set Enrichment Analysis (GSEA) was performed using the fgsea package, with genes ranked by a combined score of Log2FC x Log10(FDR). GSEA was run against four gene set databases: MSigDB Hallmark gene sets (H), MSigDB ImmuneSigDB (C7), REACTOME pathways, and Gene Ontology Biological Process (GO-BP) terms, accessed via the msigdbr and ReactomePA packages. Over-representation analysis (ORA) was performed in parallel using clusterProfiler, applying enrichGO (GO-BP) and enrichPathway (REACTOME) functions on filtered DEG lists. For ORA, GO terms were simplified using a semantic similarity cutoff of 0.6 to reduce redundancy. Multiple testing correction was applied using the Benjamini-Hochberg method, and pathways with adjusted p-value < 0.05 were considered significant Results from both GSEA and ORA were integrated for visualization, with GSEA-derived pathways displayed as dot plots colored by Normalized Enrichment Score (NES) and ORA-exclusive pathways annotated separately. For endothelial cells and fibroblasts genes analysis, genes had to be significantly different between the two sexes within at least one cluster to be selected.

Ligand-receptor mediated cell-cell interaction were inferred using CellChat (version) in R. Suture-associated fibroblasts, periosteal fibroblasts, LEC and Artery (Bmx+) clusters were excluded from this analysis due to either anatomical constraints limiting their interactions, or insufficient cell numbers. CellChat analysis was run separately on male and female datasets, and the resulting interaction networks were compared to identify sex-driven difference in interaction number, interaction probability, and ligand-receptor pair usage across dura cell population. Differential interaction probabilities were visualized using ggplot 2 custom R scripts.

To assess the potential relevance of sex-biased dural gene expression to neurological diseases with documented sex prevalence differences, sex-biased DEGs identified across all dural cell types were cross-referenced against curated disease gene sets using the DisGeNET database and the GWAS Catalog, queried programmatically in R. Ten neurological and autoimmune conditions were selected based on their known sex prevalence ratios and relevance to meningeal biology, including female-predominant conditions (Alzheimer’s disease, multiple sclerosis, lupus/SLE, migraine, and depression) and male-predominant conditions (glioblastoma, autism spectrum disorder, Parkinson’s disease, Fragile X syndrome, and bacterial meningitis). For each disease-cell type combination, the number of overlapping DEGs was quantified and classified as concordant (sex-biased direction of gene expression matching the sex prevalence of the disease), discordant, or mixed. Results were visualized as a bubble matrix in R using ggplot2, where bubble size represents the total number of overlapping disease-associated DEGs and bubble color reflects the concordance classification.

### Statistical analysis

Sample sizes were chosen based on standard power calculations using similar published experiments. Statistical methods were not used to recalculate or predetermine sample sizes. Statistical test usage is described in each figure. The numerical value of the statistical test is displayed on the graphs. Animals with biological (hydrocephaly, runt …) or experimental (poor perfusion) abnormalities were excluded from analysis. Statistical outliers were removed based on the Grubbs test with a significance level of 0.05. Choice of test were decided based on number of mice per group and distribution. Statistical analysis was performed using Prism 10 (GraphPad Software Inc).

## Acknowledgements

We thank all the members of the Louveau lab and the members of the Neuroimmunology group in the Neuroscience department of the Cleveland Clinic Research for their valuable inputs during discussions of this work.

## Contributions

Concept: AL; Investigation: NA, NMF, GAT, AV, DS. Formal analysis: NA, GAT, AV, DS. Resources: JP, DD, CB. Writing: AL

---

**Supplementary Figure 1 (associated to figure 2): Sex influences the immune composition of the dura.**

A. Representative contour plot of IgM and IgD expression by meningeal B cells between male and female mice.

B. Quantification of the number of B cells subtypes in the dura of male and female mice. Mean ± sem. Welch’s test.

C. Representative dot plot of monocytes (Ly6C+ Ly6G-) and neutrophils (Ly6C int Ly6G+) in the dura of male and female mice.

D. Quantification of the number of monocytes and neutrophils in the dura of male and female mice. Mean ± sem. Welch’s test.

E. Representative dot plot of CXCR2 expression by dural neutrophils in male and female mice.

F. Quantification of the percentage and number of CXCR2 expressing neutrophils in the dura of male and female mice. Mean ± sem. Welch’s test.

G. Representative contour plot of CCR2 expression by dural monocytes in male and female mice.

H. Quantification of the percentage and number of CCR2 expressing monocytes in the dura of male and female mice. Mean ± sem. Welch’s test.

I. Bubble plot of expression of flow cytometry markers to identify meningeal macrophages clusters.

J. Quantification of the percentage of different macrophage subtypes in the meninges of male and female mice. Mean ± sem. Welch’s test.

K. Representative contour plots of T lymphocytes in the dura of male and female mice.

L. Quantification of the number of dural lymphocytes subtype between male and female mice. Mean ± sem. Welch’s test.

M. Quantification of the ratio of dural CD4 to CD8 T cells between male and female mice. Mean ± sem. Welch’s test.

**Supplementary Table 1:** List of DEG between male and female dural cells, immune cells, vascular cells, fibroblasts and mural cells.

**Supplementary Table 2:** List of DEG for immune cell clusters. Pathway analysis for meningeal ILC2s.

**Supplementary Table 3:** List of DEG for dural endothelial cells cells.

**Supplementary Table 4:** List of DEG for dural fibroblasts.

**Supplementary Table 5:** CellChat interaction analysis.

**Supplementary Table 6:** Disease gene analysis.

