## Supplementary figures and images for "A Single-Cell Atlas of the Mouse Dural Meninges Reveals Pervasive Sex Differences Across Cellular Compartments"

### Supplementary Figure 1

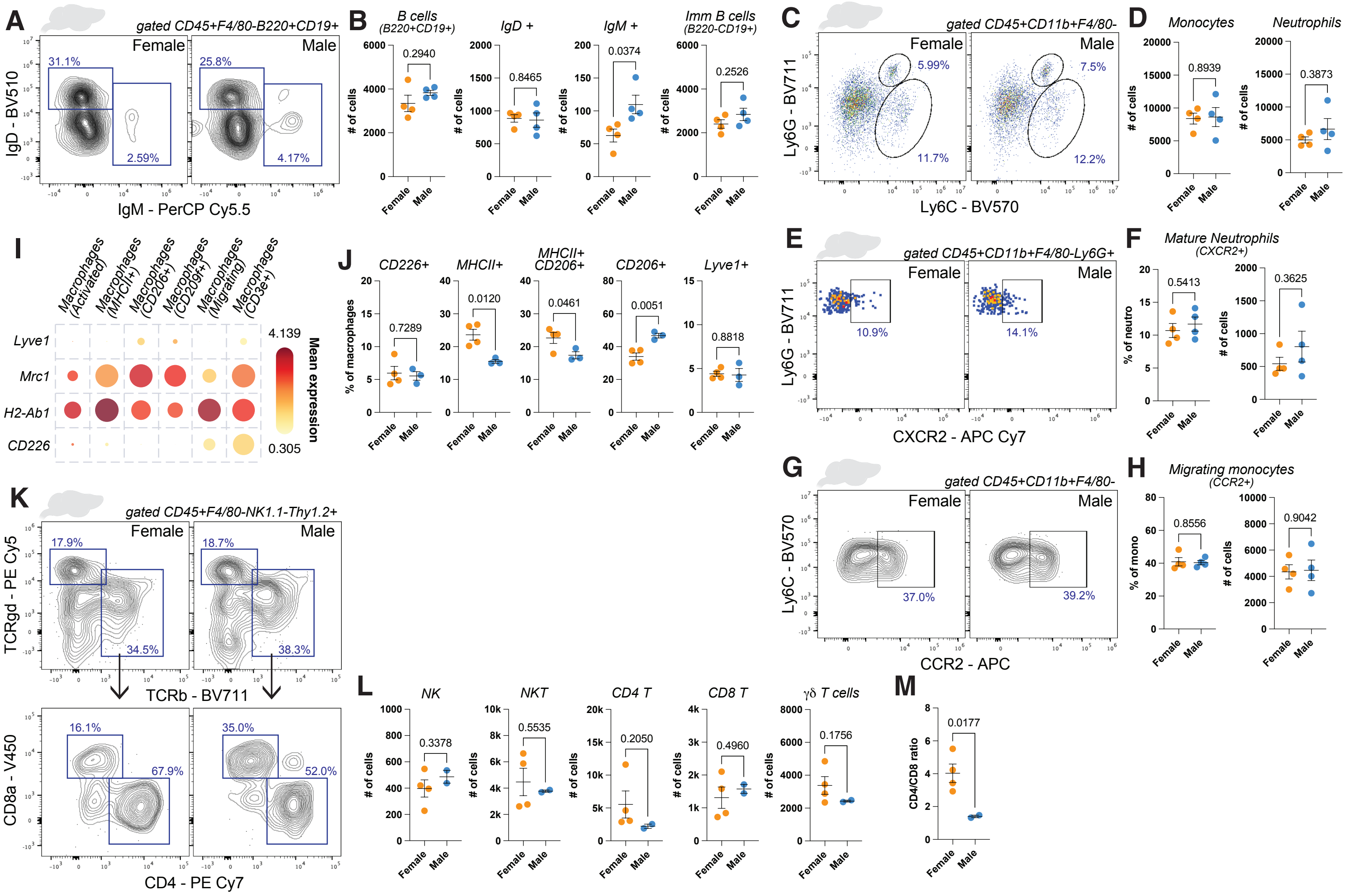
